# Policy-relevant indicators for invasive alien species assessment and reporting

**DOI:** 10.1101/2021.08.26.457851

**Authors:** Melodie A. McGeoch, Eduardo Arlé, Jonathan Belmaker, Yehezkel Buba, David A. Clarke, Franz Essl, Emili García-Berthou, Quentin Groom, Marie V. Henriksen, Walter Jetz, Ingolf Kühn, Bernd Lenzner, Carsten Meyer, Shyama Pagad, Arman Pili, Mariona Roigé, Hanno Seebens, Reid Tingley, Joana R. Vicente, John R.U. Wilson, Marten Winter

**Author notes:** Corresponding author: Melodie A. McGeoch. **Email:**.

## Abstract

Invasive alien species are repeatedly shown to be amongst the top threats to biodiversity globally. Robust indicators for measuring the status and trends of biological invasions are lacking, but essential for monitoring biological invasions and the effectiveness of interventions. Here, we formulate and demonstrate three such indicators that capture the key dimensions of species invasions, each a significant and necessary advance to inform invasive alien species policy targets: 1) Rate of Invasive Alien Species Spread, which provides modelled rates of ongoing introductions of species based on invasion discovery and reporting. 2) Impact Risk, that estimates invasive alien species impacts on the environment in space and time and provides a basis for nationally targeted prioritization of where best to invest in management efforts. 3) Status Information on invasive alien species, that tracks improvement in the essential dimensions of information needed to guide relevant policy and data collection and in support of assessing invasive alien species spread and impact. We show how proximal, model-informed status and trend indicators on invasive alien species can provide more effective global (and national) reporting on biological invasions, and how countries can contribute to supporting these indicators.

## Introduction

Biological invasions are among the most impactful drivers of biodiversity change (1, 2), in many cases with significant ecological and socio-economic impacts (3, 4). Yet there is a lack of comprehensive and readily accessible knowledge about fundamental dimensions of biological invasions, such as the rate of new species arrivals, the impacts they cause, and knowledge about their occurrences (5, 6). Nonetheless, the last half decade has seen a steep increase in the collation, harmonization, and mobilization of data on invasive alien species (IAS), e.g. (7, 8). These first extensive analyses of biological invasions show that the documented numbers of IAS have continued to increase over recent decades, with particularly steep rises on islands and in coastal regions (9). Evidence of the drivers and pathways of invasion have also improved significantly (10-13), and efforts are growing to assess invasion impacts on the environment (14, 15) and on human livelihoods (3, 16).

Alongside developments in invasion research, there has been growth in global demand for policy-relevant information on the status and trends in IAS (17). Multinational agreements developed for the purposes of addressing such global challenges call for information on the changing status of biological invasions, including the extent to which management efforts have succeeded (18). Such calls for policy-relevant information include, for example, goals and targets set by Parties to the Convention on Biological Diversity (CBD), those encompassed within the United Nations Sustainable Development Goals (in particular SDGs 14 and 15), the European Parliament Regulation on IAS, and most recently the IAS Assessment of the Intergovernmental Science-Policy Platform on Biodiversity and Ecosystem Services (IPBES). To achieve the mandates of these initiatives, recent advances in IAS information must be translated into robust, contemporary, and meaningful indicators of the status and trends in biological invasions.

However, current indicators, specifically those containing species information on IAS (e.g. rather than IAS driver or policy response indicators), fall short in multiple ways. Most fail to report on uncertainty and coverage is spatially and temporally inadequate (19). As a result, policy does not benefit from an adequate understanding of this inherently transboundary form of global change in the form of scientifically sound, relevant IAS indicators (1). There are a number of remaining challenges to delivering policy- and management-relevant knowledge on IAS so that efforts can be focused to address them (20, 21). Progress in mobilization of existing IAS data has outpaced the partnerships and processes needed to enable effective evidence-based policy. While collaborative progress has been made across data holders, IAS experts, and the agencies tasked with delivering scientifically sound and policy-relevant evidence (22), processes have yet to be put in place to maintain the generation of new data and to fill remaining taxonomic and geographic data gaps. Without a solid, repeatedly measured and up-to-date evidence-base, progress to prevent and reduce the negative consequences of IAS is hindered, and neither the evaluation nor the achievement of policy targets is feasible.

As introductions of IAS continue within and across countries globally (23), we see multiple, long-term demands for information on the status and trends in biological invasions as the evidence-based foundation for multilateral invasion policy. Draft Target 6 of the ongoing CBD Post-2020 Global Biodiversity Framework (GBF) negotiations stipulates, *inter alia*, achieving a 50 % reduction in the rate of new introductions, and the control of IAS to eliminate or reduce their impacts (24). “Rate of IAS spread” is proposed as the headline indicator for annual progress reporting by the Parties to the Convention (25). In draft Target 21, the Framework recognizes the need for quality information for the effective management of biodiversity. The associated draft indicator requires reporting on growth in Biodiversity Information relevant to the Framework (25), which would include information on species found outside of their native ranges that have negative impacts.

Here, we formulate and demonstrate three pertinent indicators to support these policy reporting needs: (i) rate of IAS spread, (ii) impact risk from IAS, and (iii) growth in information on the status of IAS. Using recent advances in IAS data collation and delivery (22, 26) and plants and amphibians as demonstration cases, we show how the indicators capture complementary facets of information, deliver interpretable and comparable evidence of trends in biological invasions at global and national scales, and simultaneously support efforts to improve relevant data coverage for future reporting. We outline the key dimensions of the data, methods and processes essential to informing on IAS. Significant recent momentum in biodiversity informatics in support of IAS data (26-29), and guidance from Essential Biodiversity Variables for species populations (30), together provide a roadmap for linking invasion targets with primary data needs and indicators (20). We foresee a step-change in indicator delivery for this key feature of undesirable environmental change.

### Policy-relevant indicators in the Essential Biodiversity Variable framework

Essential Biodiversity Variables (EBVs) encompass the core set of measurements essential for tracking biodiversity change, including IAS (20, 31). They lie at the centre of data-to-decision workflows for biodiversity policy reporting, supported by a globally coordinated network (32). The conceptual and infrastructure framework provided by EBVs works to promote strategically targeted data collection, and standardized data integration and modelling to deliver the biodiversity information needs of research, policy and management (33).

Here we identify and develop a complementary set of globally and regionally applicable indicators drawing on the EBV-centred framework that has been proposed to sustainably deliver knowledge on IAS (20). This includes in particular the Species Distributions EBV, i.e. the probability of occurrence for the global extent of a species group over contiguous spatial and temporal units (20, 33), as a basis for estimating trends and uncertainty in IAS range expansion (spread) (see *SI Appendix*, Fig. S1 for operational definition of IAS). Added to this is information on species traits and ecosystems informing on IAS impacts. While the full realization of the delivery of IAS indicators from an EBV framework has some way to go, key steps towards linking invasion policy targets with the data and models needed to support ongoing policy tracking at global and country scales are being made (22, 26, 28). Below we outline three indicators derived from spatially and temporally explicit data on alien species and their impacts as a necessary contribution to realizing this vision (19) (Table 1).

**Table 1.** The purpose and interpretation of indicators of invasive alien species spread, impact and information growth at global and national scales. Important parts of the process of identifying and adopting indicators for reporting on policy and management effectiveness are (i) establishing clear links between general and specific goals and targets, globally and by individual countries, and (ii) the making explicit how the indicator can be used to evaluate success; articulated below for the three indicators.

|  | Indicator | Description | Policy Goal | How will we know when the goal is achieved? |
| --- | --- | --- | --- | --- |
| 1 | Rate of Spread | Trends in numbers of invasive alien species and the rate of spread estimated from this trend, at global or country scales | To slow the spread of invasive alien species to achieve a reduction in the rate of new introductions (e.g. in support of Target 5 of the Post-2020 GBF) | The estimated change in IAS introductions is negative and declining in the country or region, brought about by the prevention of new introductions and the eradication of species already established. |
| 2 | Impact Risk | Trends and distributions of impact risk from invasive alien species, at global and country scales | Eliminate or reduce the growth in impact risk from invasive alien species (e.g. in support of Target 5 of the Post-2020 GBF), brought about by preventing new introductions and the control and eradication of established species | The growth and spread of those IAS with the severe, or priority impacts and types of impacts, is halted and reversed through global, regional and local prevention, eradication, population control and containment actions. |
| 3 | Status Information | Growth in invasive alien species information, at global or country scales | Improve the status of policy-relevant invasive alien species information available to researchers, decision-makers and the public (e.g. in support of Target 19 of the Post-2020 GBF) | All countries have information covering the introduction, impact and spread of all established IAS in the country, achieved *inter alia* by actions outlined in Table 2. |

#### I. Rate of Invasive Alien Species (IAS) Spread

##### Definition

*The change in number of invasive alien species over time that are expected to have established in a new region (i*.*e*., *rate of new introductions) based on observed trends in IAS observations*. This indicator can be disaggregated by taxon, region, country, pathways or high priority sites such as protected areas.

A time series demonstrating change in the number of IAS in a country, region or globally is one of the most intuitive ways to visualize the growth in biological invasions. Indeed, trends in alien species numbers (expressed either as number per time or as cumulative alien species numbers) are one of the most frequently referred to indicators of biological invasions (9, 10, 34, 35). High and increasing introduction rates are reason for concern, suggesting that prevention and control measures have been ineffective. By contrast, from a policy perspective, low or declining introduction rates suggest that prevention efforts are succeeding or adequate. However, as we show below, raw time series of IAS discoveries are fundamentally misleading and can result in misinformed policy.

The observed trend in the number of IAS over time is a function of the rate of introduction and the rate of discovery of new species populations. An introduction rate naively estimated from the cumulative numbers of introductions over time does not take into account the sampling process that is well-known to bias observed numbers of IAS (36, 37). An estimate of genuine trends in IAS introductions therefore should include an estimation of the discovery rate. Discovery probability tends to vary over time as a result of properties of the IAS population, e.g., increase in abundance post introduction; and properties of survey design, such as sampling effort and approaches applied over time and across habitats, as well as in documenting and reporting the detection (38-41). Even when the underlying introduction rate is constant, trends in sampling effort and reporting (‘sampling effects’) can result in strong trends in new species detections (42). Further, a constant sampling effort can also result in accelerating rates of new alien species discoveries without any change in true introduction rates, in the presence of imperfect detection, periodic concerted survey efforts, and recording lags (many species are detected later than they are introduced (36, 37)). It is, therefore, critical to take the species discovery process explicitly into account when estimating trends in alien and invasive species richness.

The rate of IAS spread indicator demonstrated here therefore differentiates between the effects of species discovery and species occurrence and is based on deduced introduction and establishment rates by incorporating sampling effects. IAS richness is modelled to account for both the introduction and sampling processes (36) to provide an improved estimate of the underlying introduction rates. The simplest such model (36), referred to here as the “sampling model”, is based only on a time series describing the number of detected species in a given time period and has the advantage of not requiring data in addition to those commonly available (more complex approaches are possible, e.g. (43)). Once the model has been applied, the indicator is the parameter of the model describing the (exponential) change in the underlying introduction rate over time (*β*_*1*_) (see Methods). Positive values indicate accelerating introduction rates while negative values indicate decelerating rates.

### Demonstration

We applied the sampling model to a global dataset of country-level first records of invasive alien plants (see Methods). The model of the cumulative number of invasive alien plants at new locations between the years 1800 and 2000 fits the empirical data well (Fig. 1a). However, the comparison of the estimated discovery rates (new species records per year, 1960-2000) using the sampling model with the estimated underlying introduction rates shows clearly that accounting for sampling can substantially influence the inferred patterns (Fig. 1b). For example, this analysis suggests that the introduction rate of plants globally in 2000 is likely to be lower than that estimated from observed discovery rates. Conversely, in 1960 the inferred introduction rates were higher than those estimated from discovery rates, illustrating the importance of accounting for sampling before inferring IAS richness trends. In Fig. 1c, the results of the sampling model fitted to continental trends are compared for three countries for the period 1960-2000. The estimated introduction rates show divergent patterns: Australia has an accelerating introduction rate of invasive alien plants (*β*_1_ = 0.02 ± 0.003), while the rate in the United Kingdom is decelerating (*β*_1_ = −0.02 ± 0.004). Germany shows an overall stable yet mildly decreasing trend (*β*_1_ = −0.09 ± 0.032).

**Figure 1.**
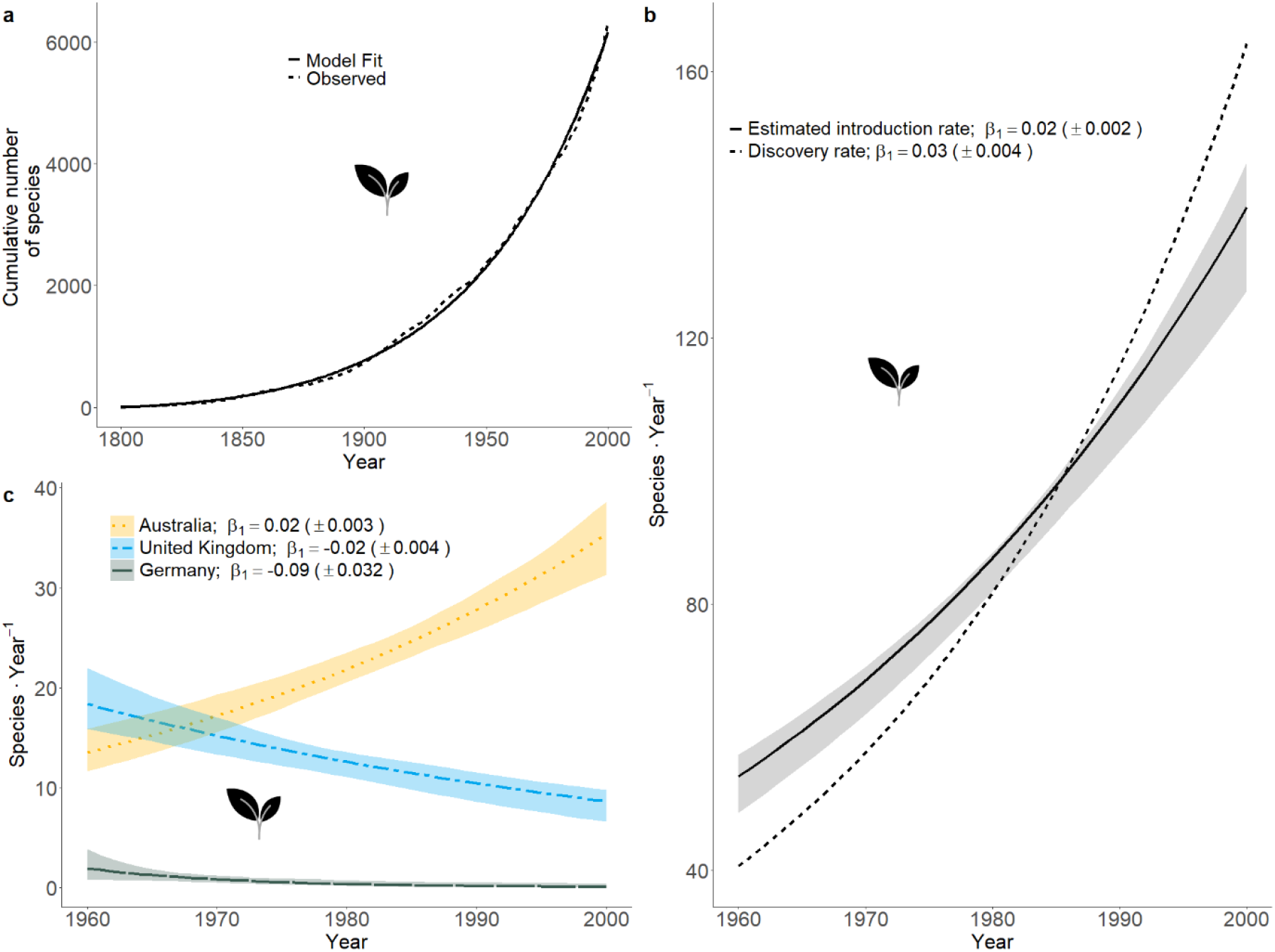
The indicator of Invasive Alien Species Spread, demonstrated using trends in invasive alien plant introductions. (a) Observed global accumulation of new invasive alien vascular plant species records (records of species in new localities) since 1800 (dashed line) and the fit of the Solow and Costello (36) model (solid line). (b) Estimated yearly introduction rate (new species records · year^-1^) in the years 1960-2000, without accounting for sampling (dashed line), and after accounting for sampling (solid line, shaded area denotes 95% confidence intervals). The annual change in rate parameter *β*_1_ represents the indicator “Rate of IAS spread” (c) The Rate of IAS Spread indicator (*β*_*1*_) calculated separately for Australia (orange; dotted), United Kingdom (blue; dash-dotted), and Germany (grey, dashed), along with the respective 95% confidence intervals.

While model-based correction for sampling effects has clear advantages, its usefulness will depend on the fit between model assumptions and reality, as with all models. Specifically, the model assumes that the underlying introduction rate changes exponentially (increasing or decreasing) with time. In addition, it assumes that sampling effort changes monotonically with time. Neither assumption holds across the board (Table 2). Deviations from these assumptions are hard to estimate, but may change the estimated underlying introduction rate. More complex models are available to account for sampling effects, such as those that rely on independent data to estimate sampling effort, e.g. (37). However, currently all models need rigorous quantification of their performance and sensitivity to deviation from model assumptions.

**Table 2.**
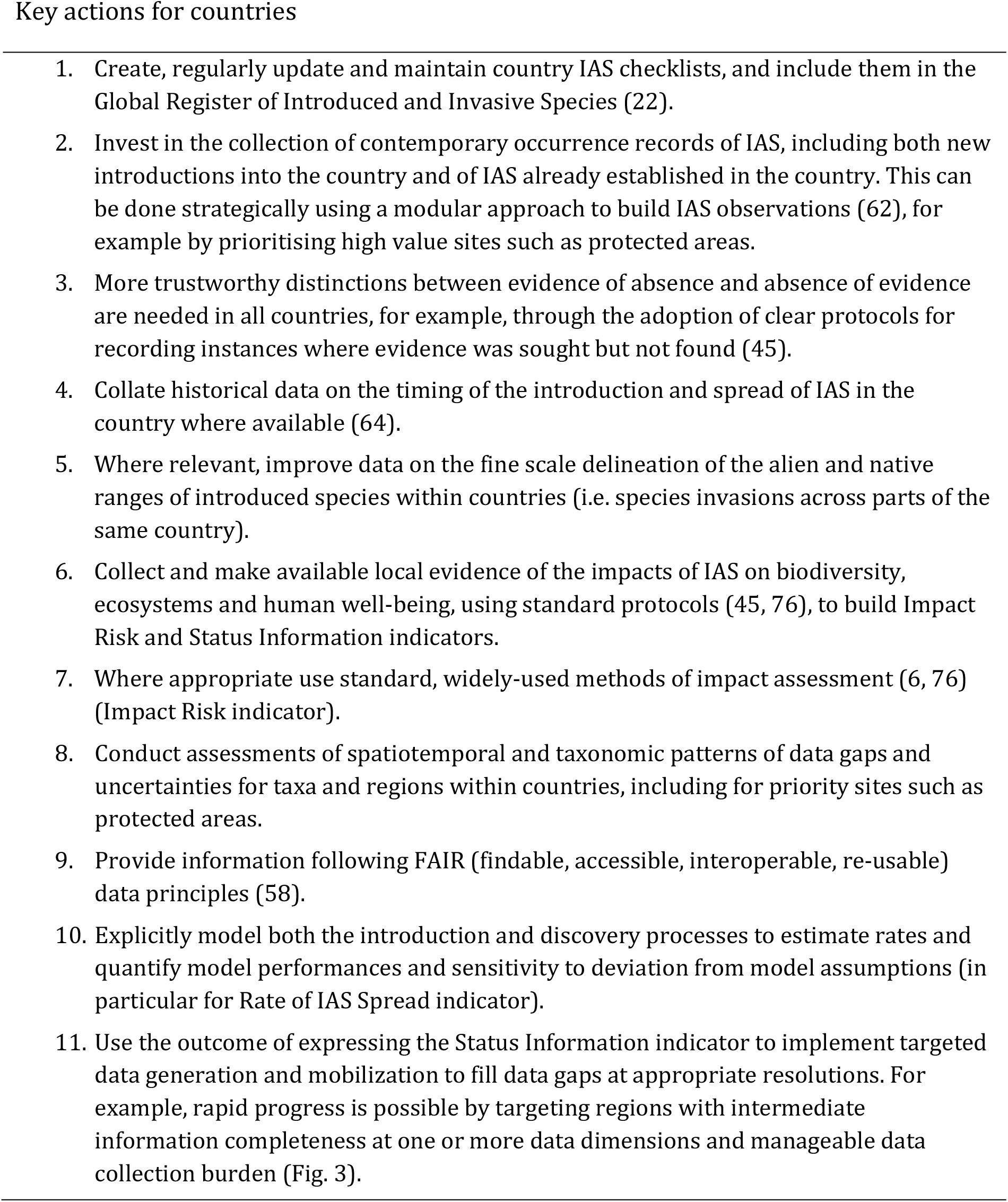
How countries can contribute to invasive alien species indicators. Key actions countries can take to enable the use and increase the relevance of these indicators at a national scale, and to simultaneously improve the information value of these indicators for global reporting.

Another factor to consider when using modelled estimates of invasion trends as an indicator is the time period used to calculate the indicator. Time series dating back many decades are suitable for estimating long-term trends, but are less relevant for assessing the effectiveness of contemporary measures to reduce the rate of new invasions. We note that the length and recency of the time series over which rates are computed may impact invasion trends. Although invasion trends are often expressed dating back to the 1800s, the period from ∼1970 is more appropriate for reporting on the effectiveness of invasion policy, *sensu* (44) (Fig. 1). Ongoing growth in data, e.g. from citizen science contributions, combined with workflows and technology for a timely and geographically-specific capture of observations offers new opportunities for measuring observed spread (19, 32), and for reporting and visualizing species-specific and combined trends in invasions.

#### II. Impact Risk: Trends and distributions of impact risk from invasive alien species

##### Definition

*The change in compound impact risk of invasive alien species that are expected in a region given general observation trends and available environmental impact data*. This indicator can be expressed as a trend sub-indicator or a species distribution sub-indicator, and disaggregated by taxonomic group, region, country and type of impact to prioritize impacts and sites to eliminate or reduce these impacts.

Evidence of IAS impacts on biodiversity and ecosystems must be taxonomically, spatially and temporally explicit if it is to robustly inform decision-making at appropriate scales. Currently, evidence of IAS impacts is biased towards population and community level effects and short-term impacts (6, 45). Even for the most well-studied species, ecosystem-level evidence is rare (6). Nonetheless, cumulative impacts of IAS are known to compound independently and synergistically across multiple IAS at particular points in space and time (13, 46). Being able to communicate to policy makers the growth, distribution and potential implications of the combined impact of invasions, i.e. environmental exposure to the types of impacts caused by IAS, is a key step towards the strategic insight needed to inform an adequate and effective response.

The impact of a species in its novel range accrues as an established alien population grows and spreads. However, the severity of a species’ impact results from a complex interplay between species populations, functional traits and the environment (47). Invasive alien species impact the environment in multiple ways, e.g. via predation or disease transmission, and individual IAS may have multiple types of such impacts (15). The cumulative impacts of different IAS present at the same locality, with the same or different types of impacts, will ultimately determine the consequences of invasion for local biodiversity (48, 49). Estimates of area impacted by IAS are prone to similar survey effort and detection biases outlined for the rate of IAS spread (above). In addition, evidence is needed on (i) realized impact (not all alien species cause harm, or the same harm, in all parts of their introduced range and impact severity can also vary with time since introduction) (5), (ii) the types of impact involved (some impacts have more severe and immediate consequences than others, and different types of impact can require different management approaches), and (iii) given the large numbers of alien species now present in many localities, impact evidence is needed for full local assemblages of IAS (9) to account for compound and potentially synergistic effects, such as invasional meltdown (50). Understanding and tracking the compound consequences of biological invasion for biodiversity and ecosystems, beyond the impacts of individual species, is therefore key to informing a policy response and prioritizing investment in prevention and control.

Recent efforts to overcome the dearth of spatially-explicit, comparable measures of impact within and across taxonomic groups include the Global Register of Introduced and Invasive Species (GRIIS), which provides global and taxonomic coverage of evidence-based and nationally verified records of realized environmental impact by alien species at a country scale (22). In addition, Environmental Impact Classification of Alien Species (EICAT) assigns one or more types (mechanisms) of impact to alien species (14). Together, these resources enable early indicators of the distribution and trends in cumulative impact risk from IAS (Table 1).

### Demonstration

We demonstrate a multispecies indicator of IAS environmental impact that captures spatial and temporal dimensions of impact risk to biodiversity and ecosystems, using amphibians as the demonstration case. We define ‘impact risk’ as the presence (observed or estimated) of one or more IAS known to negatively affect biodiversity via one or multiple types of impact. The indicator aims to deliver information on the trends and distributions of impact risk from multiple species and the ways in which they impact biodiversity (Table 1). The indicator combines global and country-level evidence of impact risk with types of impact (from GRIIS (22) and EICAT assessments (51)), country-level occurrences of IAS (from GRIIS), modelled species distributions (using ecological niche models), and first records of introduction (9) to track growth in multiple impacts over time. The indicator can be expressed in two ways: (i) as a trend, measured as the cumulative number of IAS with evidence of impact by impact type over time, and (ii) as the spatial distribution of impact risk derived by summing spatially-explicit relative likelihoods of occurrence from ecological niche models for each impact type (*SI Appendix*, Supplementary text) (Fig. 2).

**Figure 2.**
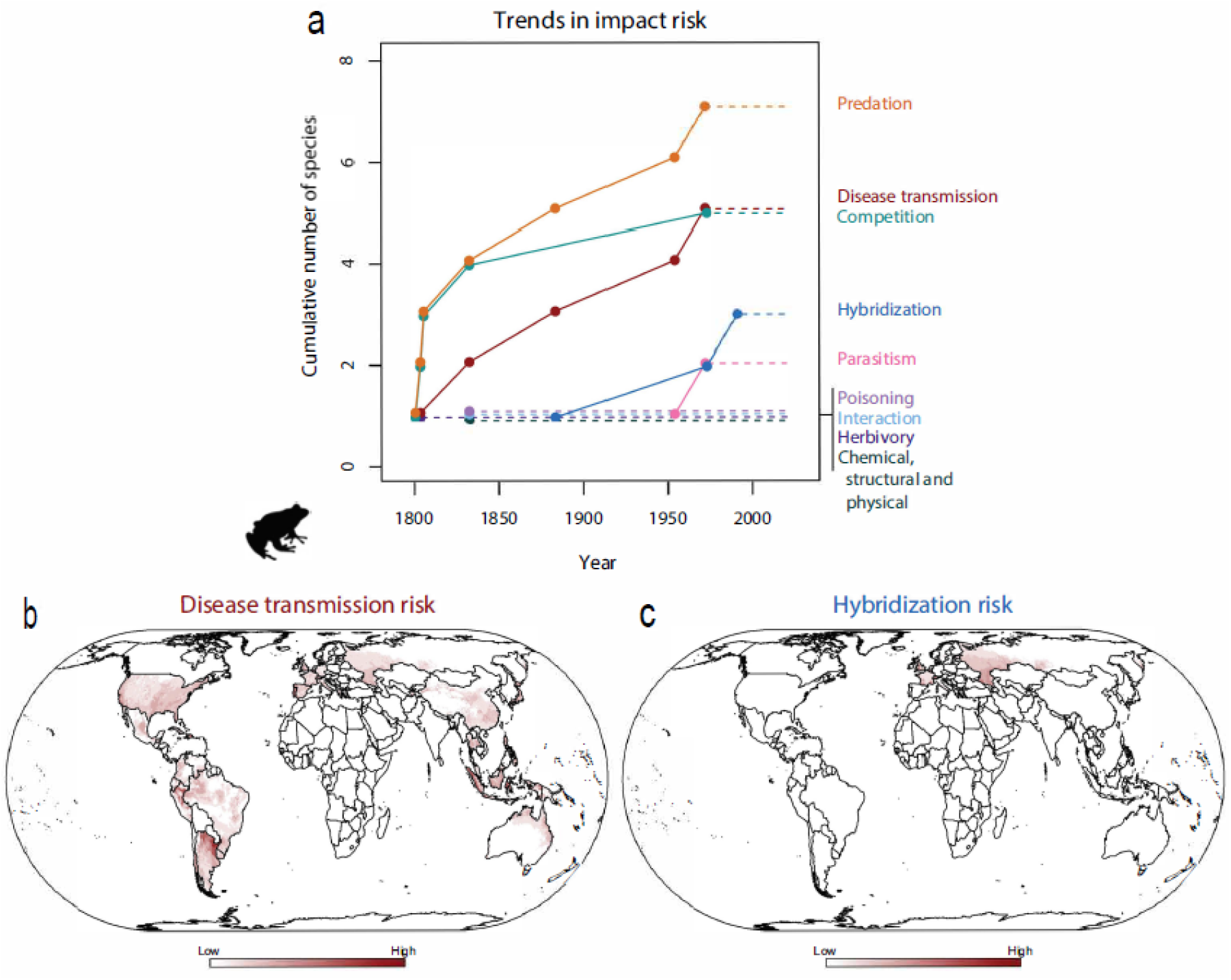
Global trends and distributions of impact risk from invasive alien amphibians. **A**. The global growth in impact risk to biodiversity associated with amphibian invasions, represents the growth in the number of species known to be associated with nine types of impact. Current indicator status is assumed to be at the level of the most recent record of introduction (dashed lines, e.g. 7 for predation). **B-C**. Impact risk maps, illustrating the distribution of impact risk from invasive alien amphibians. **B** shows the distribution of risk from the subset of invasive alien amphibians considered to transmit disease, and **C** shows the distribution of hybridization impact risk. These distributions represent the projected global risk of impact expressed as summed global climate suitability of species across regions and in countries where these multiple species are known to have been introduced, estimated using ecological niche models (*SI Appendix*, Supplementary text). In each case the sums are based on the relevant species and subset of types of impact of concern, among the total 11 species and 9 impact types (*SI Appendix*, Supplementary text).

The trends in IAS impact risk show, for example, that global growth in impact risk from amphibians is dominated by predatory species, followed by those that transmit diseases or have a high competitive ability, while comparatively few amphibian IAS carry a hybridization risk (Fig. 2a). Estimated disease transmission risk is widely distributed globally (Fig. 2b), whereas hybridisation risk currently occurs predominantly across parts of Europe, Eastern Europe and Russia (Fig. 2c). It is conceivable that this indicator could show a reduction in impact risk over time with successful local eradication and control efforts resulting in a decline in the number of species associated with a particular type of impact risk (Fig. 2a), along with a decline in the distribution of the impact risk they pose (Fig. 2b).

Any impact risk indicators will be biased by insufficient data on regional occurrences of alien species and their impacts, which are both notoriously difficult to estimate and generally subject to time lags, e.g. (52, 53) (Table 2). The approach used here also rests on a number of key assumptions (e.g., niche conservatism; unbiased sampling effort in environmental space; equilibrium with the environment - which is notoriously violated for alien species; and that sums of relative likelihoods of occurrence approximate species richness (54, 55) (*SI Appendix*, Supplementary text). The Impact Risk indicator should, therefore, be considered alongside the Status Information indicator (below) to assess gaps in information and their consequences for interpretation. Future development of impact risk indicators for more inclusive sets of species and taxa could include modelled species trends over time (as for the Rate of Spread indicator) to provide a more robust assessment of temporal change in impact. Impact risk distributions can also be constructed for priority species subsets, such as those considered to have massive environmental impacts according to EICAT (14). It is likely that available impact evidence relevant for such indicators will grow, including for example regional impact evidence (countries and islands) from GRIIS (22), information on socioeconomic in addition to biodiversity impacts, e.g. (16), and more sophisticated ways of estimating compound and synergistic impacts, e.g. using network analysis (56).

#### III: Status Information: Growth in invasive alien species information

##### Definition

*Growth in the coverage of spatiotemporally explicit occurrences and impact data for invasive alien species*. This indicator can be disaggregated by taxonomic group, region or country.

An indicator of the completeness of information on the key dimensions of the status of IAS, i.e. spatiotemporal change in their distributions and environmental impacts, could leverage targeted investment in the collection of the data most needed to inform policy and management efforts (Table 1) (57). A Status Information indicator would, by highlighting the gaps, also over time lead to an improvement in the information value of other IAS indicators, in particular those on IAS spread and impact, supporting a more general biodiversity information target as currently envisioned by the Post-2020 Global Biodiversity Framework (24). An information indicator should report on the availability of data that are useful for decision-making and reporting, i.e. data that are available and accessible for this purpose and that meet a minimum set of data standards (28). Status Information should therefore be assessed using collated, harmonised and open access data (that meet FAIR principles (58)) to further ensure that indicators are repeatable and transparent, and thus can ultimately be calculated by anyone.

Quantifying data completeness for IAS is not straightforward. Simple assessments of data volume or numbers of recorded events (e.g. species, locations, dates) are not comparable because countries differ ecologically and socio-economically in multiple ways that impact data availability. Regions differ in the number of IAS populations and species present and thus in the data quantity needed to completely capture their spread and impacts (i.e., they have different ‘information burdens’). Although Essential Biodiversity Variables for species populations aim to provide modelled coverage of locality by time estimates of species distributions (33), meaningful indicators of IAS information adequacy do require baselines against which gaps in detected IAS richness or impacts can be tested (59). In addition, a systematic and repeated recording of IAS sampling effort across countries and taxa to support the rigorous control for biases is currently unrealistic, given the uneven focus and capacity of biodiversity monitoring and reporting schemes (32, 59-61). These general challenges are not dissimilar to those for native species indicators, as reflected by the current draft biodiversity information Target 20 of the Post-2020 GBF (25). Until more comprehensive data become available, however, regional (e.g. federal states, island groups) or country-wide IAS checklists (22) can offer baseline information on the alien species currently known to occur in those regions, and that are known to impact native biodiversity (IAS, *SI Appendix*, Fig. S1). Against these baselines, meaningful metrics of IAS information completeness and adequacy can already be calculated.

The IAS Status Information indicator is thus designed to inform on the status and availability of the empirical data needed on the introduction, range dynamics and impacts of IAS, and to support other IAS indicators, based on three key dimensions of IAS information (62): (i) introduction date evidence - completeness of data on the introduction year for known IAS and country of occurrence (availability of first record date per alien species known to occur there and to impact biodiversity there or elsewhere, *SI Appendix*, Fig. S1); (ii) impact evidence - completeness of country-level impact evidence for known IAS; (iii) range dynamics evidence, i.e. minimum adequacy of spatiotemporally explicit occurrence records (see Methods). These three sub-indicators of status information can be averaged to a single, composite indicator of IAS status information for general reporting (see Methods). They can also be tracked individually to enable responses to be specifically targeted at each dimension of the data needed.

### Demonstration

Readily accessible information on introduction dates, range dynamics, and impacts of amphibian IAS globally and at a country grain were used to calculate the composite IAS Status Information indicator (Fig. 3a) and its three sub-indicators (Fig. 3b, *SI Appendix*, Supplementary text). From the composite indicators, Brazil is an example of a country with comparatively complete overall evidence on amphibian IAS, including complete information for two of the three sub-indicators (*In* and *Im*, Fig. 3). At the same time, however, Brazil has a low information burden with just two IAS amphibians (Fig. 3a inset; sub-national species translocations were not included in our workflow (63)). For example, for the IAS amphibian *Lithobates catesbeianus* known to be introduced in Brazil, there is both a date of first record and a verified record of its impact in the country (Fig. 3b, impact evidence and introduction date evidence). However, a minimum adequate number of occurrence records for this species is only available for 20% of the half-decade time slices since its introduction (range dynamics evidence, Fig. 3b), yielding an overall Status Information indicator score of 73% (Fig. 3a) (*SI Appendix*, Supplementary text). Brazil does have a second invasive alien amphibian, not yet included in its GRIIS Country Checklist, as well as four within-country species translocations (63). This indicator is intended over time to bridge this disconnect that exists between research data and data available for national reporting by highlighting information in this way. The indicator could further be expanded to encompass sub-national scale aliens.

**Figure 3.**
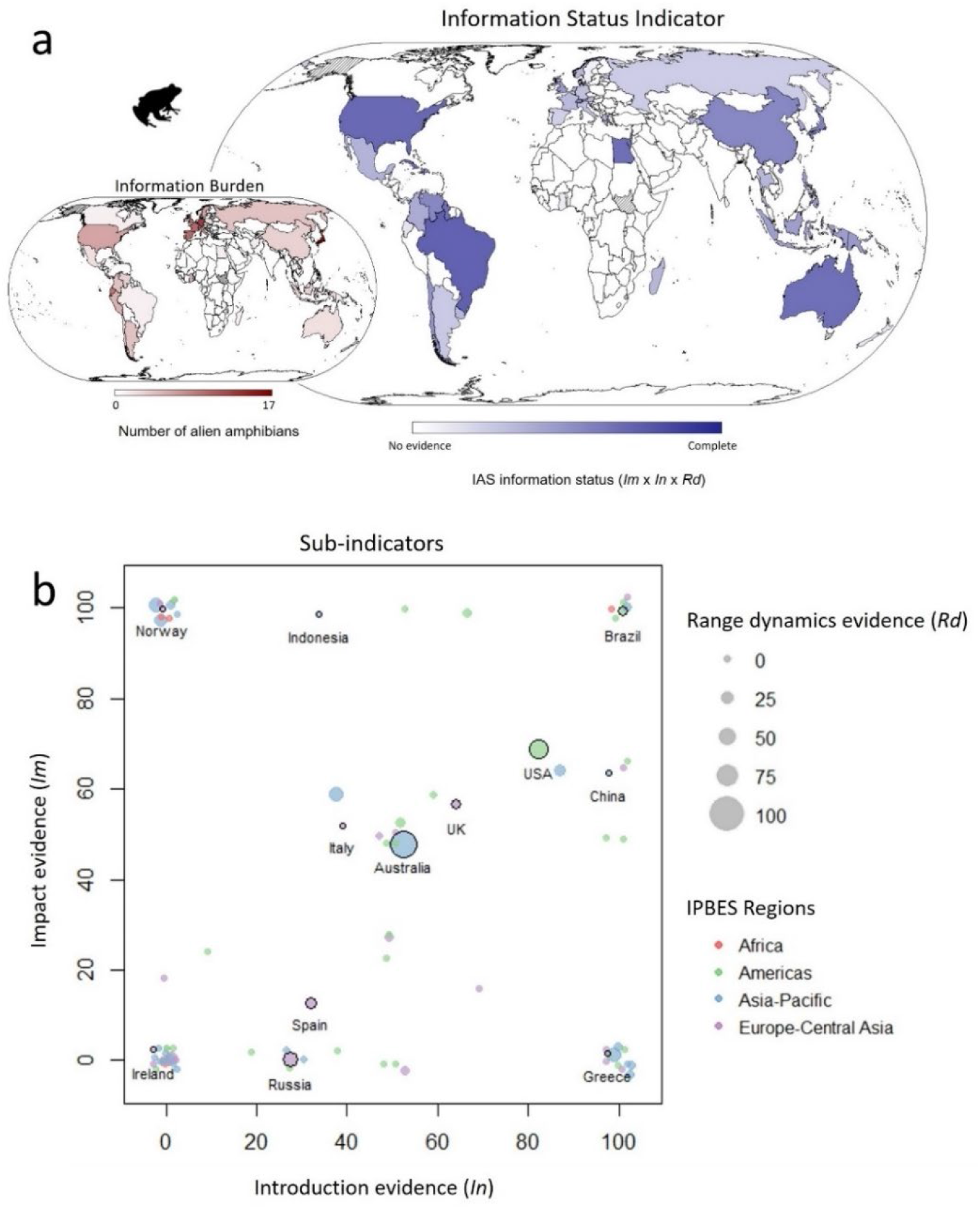
Indicator of Invasive Alien Species (IAS) Status Information. (A) Values of the indicator expressed as the average of the three sub-indicators (shown in panel B), ranging from 0 (no evidence) to 100 % (complete evidence), to be interpreted alongside the inset showing the information burden (number of known alien amphibian species) per country. Countries in white have no reported amphibian IAS, and dashed countries have no data to calculate the indicator (*SI Appendix*, Fig. S1). (B) The completeness of information on invasive alien amphibians disaggregated by sub-indicator: *Im*, percentage of IAS with national-level evidence of impact; *In*, percentage of these species for which introduction date is available; *Rd*, point size represents the percentage of species for which at least 10 occurrence records per five year period (1970 – 2019) are available. Each point is a country and selected country names are shown as examples and highlighted by in black outline. Colours show the IPBES Region that the country belongs to (74, 75).

Countries like Brazil with few amphibian IAS and thus a lower information burden could achieve 100% on each sub-indicator axis with moderate effort. By contrast, the investment needed to achieve full information coverage of the indicator is much higher in Japan, which has the most amphibians listed as IAS (17) (Fig. 3a inset). Most of Africa, the Arabian Peninsula and the Indian subcontinent have no known invasive amphibians listed (Fig. 3a inset), while Alaska and South Sudan have no data (following Fig. S1). Many countries have complete information for one sub-indicator only, including a cluster of countries with introduction evidence but little to no impact evidence (e.g., Greece, Indonesia), and another cluster (including Norway) has the opposite (Fig. 3b). The U.S.A. is one of few countries showing moderately complete information across all three dimensions (Fig. 3b; larger circles in the center and upper-right corner). Importantly, this indicator is intended for tracking information growth within countries over time, and overall at a global scale, and not for the purpose of comparing the efforts of particular countries, especially given the uneven distribution of the information burden across them (Fig. 3a inset).

The Status Information indicator necessarily relies heavily on the key data sources it draws on (see Methods). While these data sources represent recent and significant advances in data collation and accessibility for IAS, they are themselves incomplete and dynamic. A key role of this indicator is therefore to illustrate not only the need for new data on IAS to fill gaps and to track the ongoing spread of IAS, but also to highlight the importance of sustained investment in generating, improving and maintaining such data as essential for providing the evidence needed for policy and management for IAS. These data initiatives for IAS, such as GRIIS, are to date largely supported by primary research and volunteer efforts (e.g. (22, 64)). The mechanism that GRIIS provides for countries to update their IAS checklists is key to sustaining these indicators (Table 2).

### The way forward for a new age of invasive alien species indicators

We envisage a step change in reporting on biological invasions, enabled by information rich, relevant and sustainable invasion indicators. These indicators are supported by representative data, models of spatiotemporal dynamics using Essential Biodiversity Variables for species populations and global coverage at national resolution (20, 33). They are further designed to meet the properties required of scientifically valid and policy-relevant indicators, including reproducibility, uncertainty reporting, scalability across global and country reporting and encompassing spatially and temporally explicit information (19). Expressing all three indicators for multiple vertebrate groups over the post-2020 reporting cycle is feasible. For plants there is an urgent need to conduct comparative impact assessments, and such assessments for insects are under way (45). Decision-support outputs in the form of maps and indicators are already being supported by dashboards and platforms such as the GEOBON portal (https://portal.geobon.org/) and Map of Life web interface (https://mol.org/indicators), and are readily extendible to encompass the invasive alien species case.

While the indicators demonstrated here present a significant advance, a number of improvements are foreseeable with updated information sources and advances in their integration and sustainable delivery (Table 2). Perhaps most importantly, improvements in the stream of distribution data will rely on countries investing strategically in IAS monitoring (65). National-level indicators are much needed (19) and are instrumental to progress toward achieving biodiversity targets (66). The three indicators, informed by sub-national data and with ready re-scoping of the methods involved, can be quantifiable and meaningful for individual countries (Table 2). In many countries, one or more of these indicators could already be expressed at the national scale using the data resources that we drew upon here, and all three global indicators can be disaggregated to provide country estimates (e.g. Fig. 1c). While the adequacy of data and monitoring systems to populate them is variable across countries, targeted investment by countries in data on these key dimensions of invasion (6, 65) (Table 2) will provide powerful opportunities for a step change in the value of information on IAS for policy and reporting. This progress can be readily supported by open standards, FAIR data and existing intergovernmental partnerships, as well as by platforms such as Map of Life for delivering species-level reports and model-corrected estimates (67).

Ongoing investment is required not only for improving the data that are essential to reporting, but also for creating and maintaining information platforms that are able to deliver indicators at multiple scales and sustain indicator updates. As a result of such investment, a decade from now we could feasibly know the trends in occurrence, distribution and impacts of IAS, at multiple scales with measurable certainty and informed by spatiotemporally explicit data and model estimates. This information could be near real-time for priority species, and readily accessible to researchers, countries, and trading partners. Indicators of the species that threaten biodiversity, together with indicators covering drivers of invasion, the status of species and ecosystems affected, and societal responses to dealing with them, provide a robust framework for tracking and managing progress (20). Drawing on such a platform of evidence, predictions and scenario planning, e.g. (68-70) can be used to ensure that they inform decision making at a variety of spatial scales. Importantly, we envisage a pipeline of data leading to indicators that deliver the information needs of policy assessments, and biodiversity, ecosystem and sustainable development targets, as they progress across the decades.

## Materials and Methods

The dataset of taxonomic occurrences and first records was generated by integrating databases of alien species occurrences at country scales with first records of discovery using a workflow published for this purpose (26), to provide the backbone of country-level locality and introduction event data for each indicator. The workflow was used to harmonize i) taxon names according to the backbone taxonomy of the Global Biodiversity Information Facility (71), ii) the Darwin Core terminology of occurrence status and means of establishment (28), and iii) the presentation of first records in countries as single years (64). Multiple forms of uncertainty are associated with assigning alien and IAS status to populations and species (72). To ensure appropriate interpretation of indicators and their repeatability it is essential that they are underpinned by systematic decisions that operationalize the definition of species included and excluded and the use of data in the indicators (see *SI Appendix*, Fig. S1). Plants (151 countries) and amphibians (82 countries with amphibian IAS) were used as taxonomic examples to best demonstrate each indicator.

The Rate of IAS Spread indicator was produced by fitting the model described in Solow and Costello (36) to the time series of first records of IAS in each country. This model finds the maximum-likelihood (ML) estimates of five parameters used in two dependent functions, the first (*μ*_*t*_) describing the mean annual introduction rate, and the second (Π_*st*_), describing the probability of discovery of species in year *t* see (36) for full model description):

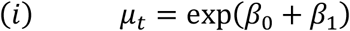

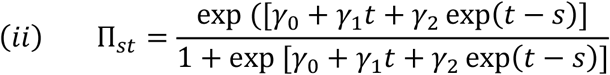

We then focused on the parameter *β*_1_ – the mean annual change in the rate of new species introductions. We used equation (*i*) to visualize the underlying introduction rates (new IAS · year^-1^ ; Fig. 1b, c). Confidence intervals were produced by resampling the parameters based on the variance and covariance of their ML estimates and calculating *μ*_*t*_ based on the resampled *β*_0_ and *β*_1_. We compared this to the discovery rate (new IAS records · year^-1^) estimated from an exponential model fit to the annual number of new IAS records.

The Impact Risk indicator is illustrated using 11 amphibian species assessed as having negative impacts on biodiversity (51), including the full suite of impact types associated with each species (*SI Appendix*, Supplementary text). The global potential impact of these species was projected based on their climatically suitable areas (full methods provided in *SI Appendix*, Supplementary text). Global projections of climatically suitable areas were based on ecological niche models calibrated using occurrence data from the species’ global ranges (native and invaded ranges, where data were available) and seven environmental variables that were deemed most relevant to amphibian ecology and biogeography. The impact risk for a species in a geographic grid cell is thus the ecological niche model estimation of the suitability of that cell for a species, summed for those IAS in the selected species set (*SI Appendix*, Fig. S1).

The species included in the Status Information indicator are those listed by GRIIS, specifically for the 72 countries where alien amphibian species are present that are associated with evidence of impact anywhere outside of their native range (Fig. S1). The Status Information indicator is composed of three equally contributing sub-indicators. (1) Introduction date evidence (*In*) is obtained by calculating the percentage of species listed by GRIIS for a given region for which there the date of first introduction is available (from (9, 64)). (2) Impact evidence (*Im*) - the percentage of species listed by GRIIS for a given region for which the impact has been assessed (i.e. designated as ‘isInvasive’ in GRIIS (22), Fig. S1). (3) Range dynamics evidence (*Rd*) - this sub-indicator relies on the availability of point record data facilitated by GBIF (73). To calculate this sub-indicator, each species-region combination was assessed as meeting a minimum criterion of having at least 10 records available for each time slice of 5 years within the 50 years considered (1970-2019). Minimum data adequacy, here, is defined as having some minimum number of occurrence records per species needed to enable modeling of IAS distributions and their change (i.e. range dynamics) (33). The minimum number is set low to make this target achievable (given adequate sampling effort) irrespective of the species’ rarity or regional population dynamics. This minimum number could readily be revised as required. We considered all unique records for each species listed, excluding duplicate records with the same coordinates and same date (day, month, and year). Only records from the period 1970 to 2019 were considered. Finally, we calculated the percentage of completeness for each region (*SI Appendix*, Dataset S1).

## Acknowledgments

sTWIST is supported by sDiv, the Synthesis Centre of iDiv (DFG FZT 118, 202548816). This work is a contribution to the Species Populations Working Group of GEO BON (https://geobon.org/ebvs/working-groups/species-populations). We thank Rachel Leihy for pre-submission comments on the manuscript. M.M. and D.C. acknowledge Australian Research Council DP20200101680. E.A., F.E., C.M. and M.W. acknowledge funding for this work through iDiv’s Flexpool mechanism. J.R.U.W. thanks the South African Department of Forestry, Fisheries and the Environment (DFFE) for funding, noting that this publication does not necessarily represent the views or opinions of DFFE or its employees. E.G.B. was financially supported by the Spanish Ministry of Science (projects RED2018-102571-T and PID2019-103936GB-C21). H.S. acknowledges funding through the 2017-2018 Belmont Forum and BiodivERsA joint call for research proposals, under the BiodivScen ERA-Net COFUND programme, and with the funding organisations German Federal Ministry of Education and Research (BMBF; grant 16LC1807A). W.J. acknowledges funding from the E.O. Wilson Biodiversity Foundation and National Aeronautics and Space Administration grants 80NSSC17K0282 and 80NSSC18K0435.

